# Bayesian Networks established functional differences between breast cancer subtypes

**DOI:** 10.1101/319384

**Authors:** Lucía Trilla-Fuertes, Andrea Zapater-Moros, Angelo Gámez-Pozo, Jorge M Arevalillo, Guillermo Prado-Vázquez, Mariana Díaz-Almirón, María Ferrer-Gómez, Rocío López-Vacas, Hilario Navarro, Enrique Espinosa, Paloma Maín, Juan Ángel Fresno Vara

## Abstract

Breast cancer is a heterogeneous disease. In clinical practice, tumors are classified as hormonal receptor positive, Her2 positive and triple negative tumors. In previous works, our group defined a new hormonal receptor positive subgroup, the TN-like subtype, which has a prognosis and a molecular profile more similar to triple negative tumors. In this study, proteomics and Bayesian networks were used to characterize protein relationships in 106 breast tumor samples. Components obtained by these methods had a clear functional structure. The analysis of these components suggested differences in processes such as metastasis or proliferation between breast cancer subtypes, including our new subtype TN-like. In addition, one of the components, mainly related with metastasis, had prognostic value in this cohort. Functional approaches allow to build hypotheses about regulatory mechanisms and to establish new relationships among proteins in the breast cancer context.

**Author Summary:** Breast cancer classification in the clinical practice is defined by three biomarkers (estrogen receptor, progesterone receptor and HER2) into hormone receptor positive, HER2+ and triple negative breast cancer (TNBC). Our group recently described a new ER+ subtype with molecular characteristics and prognosis similar to TNBC. In this study we propose a mathematical method, the Bayesian networks, as a useful tool to study protein interactions and differential biological processes in breast cancer subtypes, characterizing differences in relevant processes such as proliferation or metastasis and associated them with patient prognosis.

## Introduction

Breast cancer is one of the most prevalent cancers in the world [1]. In clinical practice, breast cancer is classified according to the expression of hormonal receptors (estrogen or progesterone) and Her2, into positive hormonal receptor (ER+), HER2+ and triple negative (TNBC). In previous studies, our group defined a new ER+ molecular subgroup, named TN-like, with a molecular profile and a prognosis more similar to TNBC tumors [2]. The remaining ER+ tumors were considered as ER-true. We also found significant molecular differences among breast cancer subtypes. For instance, differences related with metabolism of glucose were described between ER-true, TN-like and TNBC tumors [2, 3].

Proteomics provides useful information about biological process effectors and may quantify thousands of proteins. Undirected probabilistic graphical models (PGM), based on a Bayesian approach, allow characterizing differences between tumor samples at functional level [2–5]. In this study we explored the utility of Bayesian networks in the molecular characterization of breast cancer. The main feature of targeted Bayesian networks is that they provide a hierarchical structure and targeted relationships between proteins.

In this work, we used proteomics and Bayesian networks to characterize protein relationships in a cohort of breast cancer tumor samples. These networks maintain a functional structure and it is possible to use this information to build prognostic signatures. This approach also reflected previously described interactions and it could be used to propose new hypotheses and mechanisms of regulation of these proteins.

## Results

### Patient characteristics

Clinical characteristics of this patient cohort have been previously described [2, 3, 6]. Briefly, one hundred and six patients were enrolled into the study. They all had node positive disease, Her2 negative and all had received adjuvant chemotherapy and hormonal therapy in the case of ER+ tumors. Among ER+ tumors, 50 patients were characterized as ER-true and 21 were defined as TN-like (Sup Table 1) [2].

### MS analysis

Proteomics analyses from these samples has been previously described [3]. In summary, one hundred and two FFPE samples had enough protein to perform the MS analyses. After MS workflow, 96 samples provided useful protein expression data. After quality criteria, 1,095 proteins presented at least two unique peptides and detectable expression in at least 75% of the samples in at least one type of sample (either ER+ or TNBC).

### Directed networks

Using proteomics data, directed acyclic graphs (DAG) were performed. Altogether, it was possible to establish 536 edges of which 414 are guided and 122 are undetermined. These edges formed 377 components formed by different number of nodes or proteins. An overview of the number of nodes (proteins) included in each component is provided in table 1.

**Table 1:** Characteristics of the components obtained from DAG. Number of nodes = number of proteins contained in each component, Number of components = directed components obtained.

| Number of nodes | 1 | 2 | 3 | 4 | 5 | 6 | 7 | 8 | 9 | 10 | 11 | 13 | 14 | 19 | 21 | 26 | 27 | 34 | 36 |
| --- | --- | --- | --- | --- | --- | --- | --- | --- | --- | --- | --- | --- | --- | --- | --- | --- | --- | --- | --- |
| Number of components | 229 | 70 | 24 | 12 | 15 | 2 | 6 | 3 | 4 | 2 | 1 | 2 | 1 | 1 | 1 | 1 | 1 | 1 | 1 |

We characterized components from DAG analysis. Components including less than 9 nodes were dismissed because they were little informative. All components were named with the number of nodes included by the DAG analysis, and the information was completed with protein-protein interactions (PPI) based on experimental evidence obtained from Genes2FANS (G2F) [7]. For example, component 14 includes 19 nodes, 14 nodes defined from our Bayesian analysis and 5 extra nodes added by G2F.

Afterwards, components were interrogated for biological function. For components with less than 27 nodes provided by DAG analysis, this functional analysis was performed by bibliographical review. Thus, the main biological function of component 14 is metabolism. Regarding components which had more than 27 nodes provided by the DAG analysis, gene ontology analyses were done once they were completed by G2F information. Characteristics about all components are supplied in table 2 and Supplementary File 1.

**Table 2:** Features of components obtained by DAG and G2F analysis.

| Component | Main function | Total number of nodes |
| --- | --- | --- |
| Component 36 | Extracellular exosome | 65 nodes |
| Component 34 | RNA processing and mTOR pathway | 61 nodes |
| Component 19 | Proliferation | 26 nodes |
| Component 21 | Proliferation & metastasis | 34 nodes |
| Component 27 | Metastasis | 34 nodes |
| Component 14 | Metabolism | 19 nodes |
| Component 13 | Proteasome | 19 nodes |
| Component 11 | None main function assigned | 14 nodes |
| Component 10b | Proliferation | 19 nodes |
| Component 9a | Growth | 15 nodes |
| Component 9b | Proliferation & metastasis | 22 nodes |
| Component 9c | Glycolysis | 12 nodes |
| Component 9d | Transcription & ribosomes | 20 nodes |

### Component activity measurements

Component activities were calculated for each node. There were significant differences between ER-true, TN-like and TNBC tumors in the component activity for component 34: mRNA processing and mTOR, component 36: extracellular exosome, component 21: proliferation and metastasis, component 27: metastasis, component 9a: growth, component 9b: proliferation and metastasis, component 9c: glycolysis, component 10b: metastasis, component 13: proteasome and component 9d: transcription (Fig 1).

**Fig 1:**
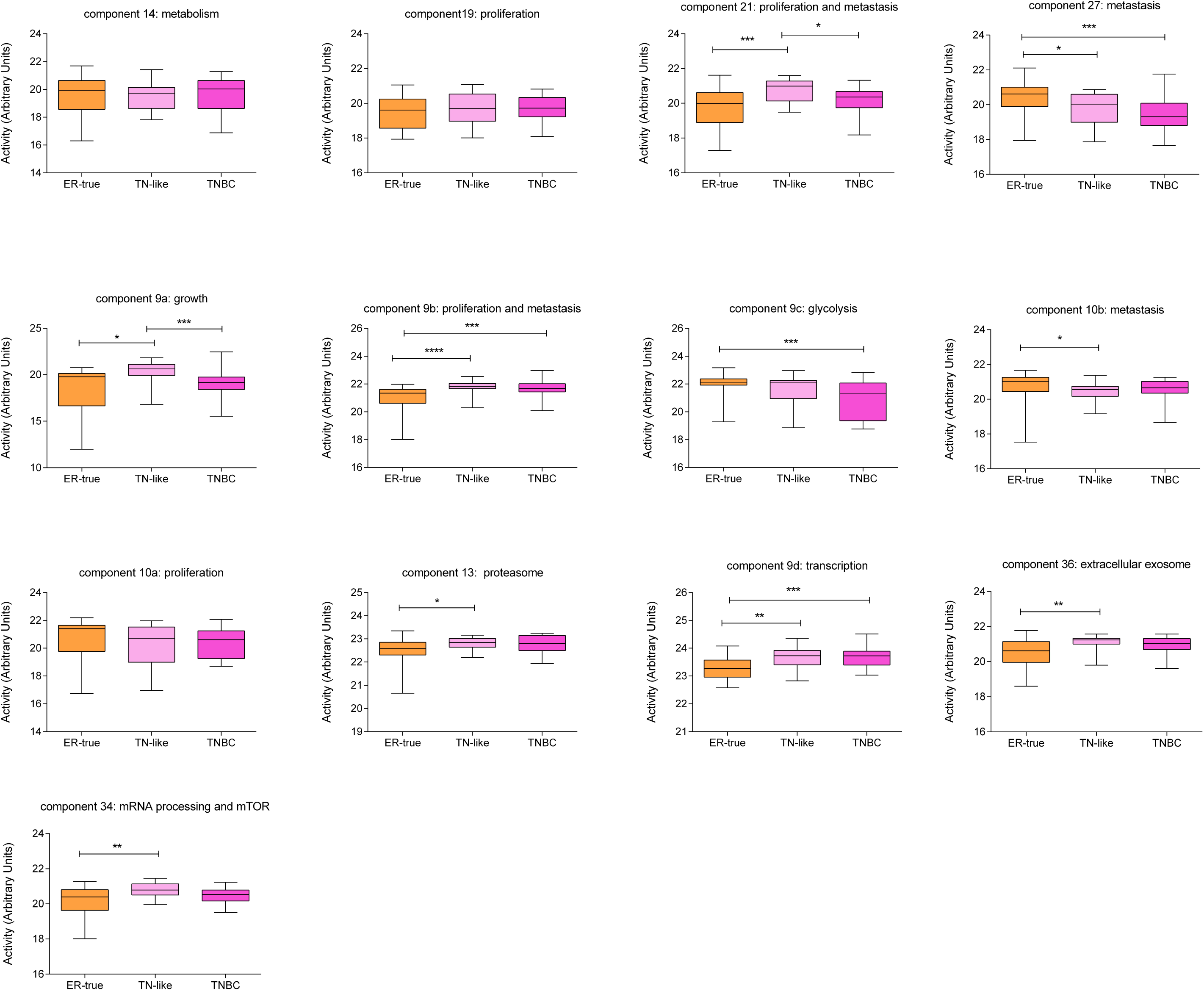
Component activity measurements for ER-true, TN-like and TNBC respectively.

### Component 10b: metastasis

Component 10b activity showed prognostic value in our series, splitting our population into a high and a low risk group and it can be used to make a distant metastasis-free survival (DMFS) predictor (p=0.0068, HR= 0.2854, cut-off 40:60) (Fig 2). Additionally, dividing by molecular subtype, this prognostic signature also split the population into low and high risk groups although it is not statistically significant (Fig 3).

**Fig 2:**
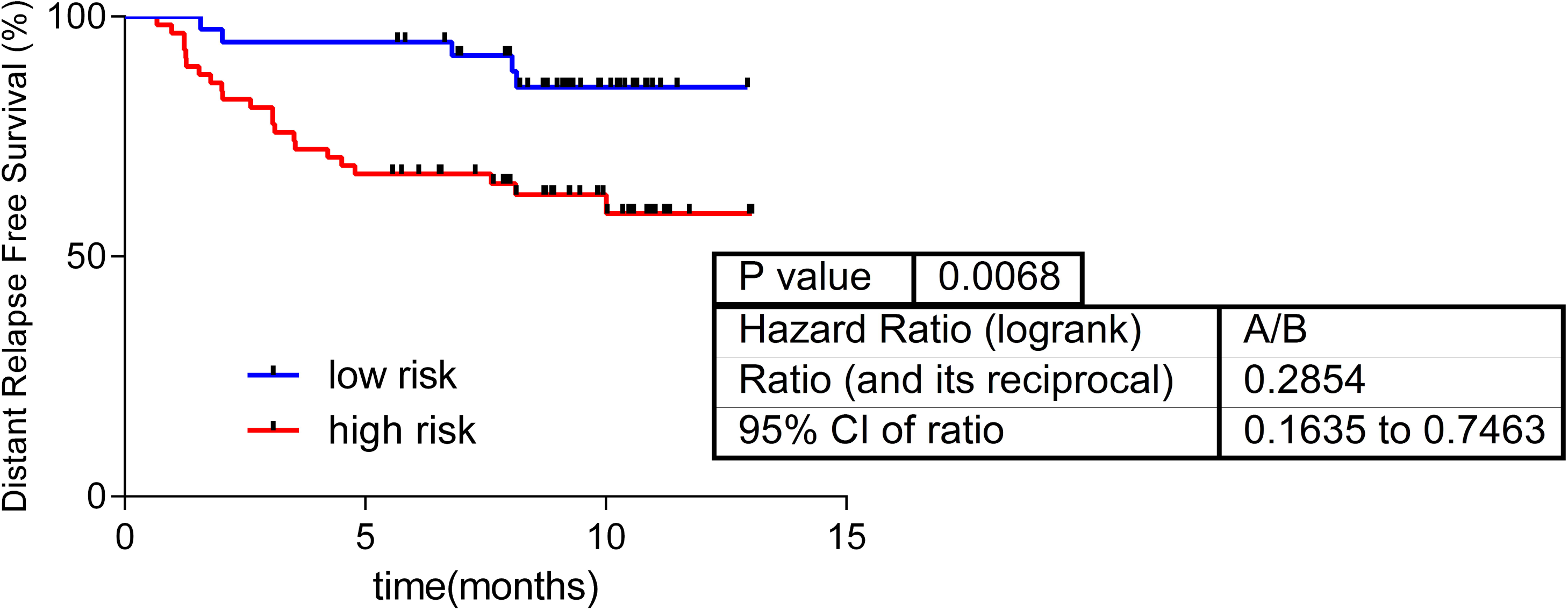
Component 10b activity prognostic value in the whole cohort.

**Fig 3:**
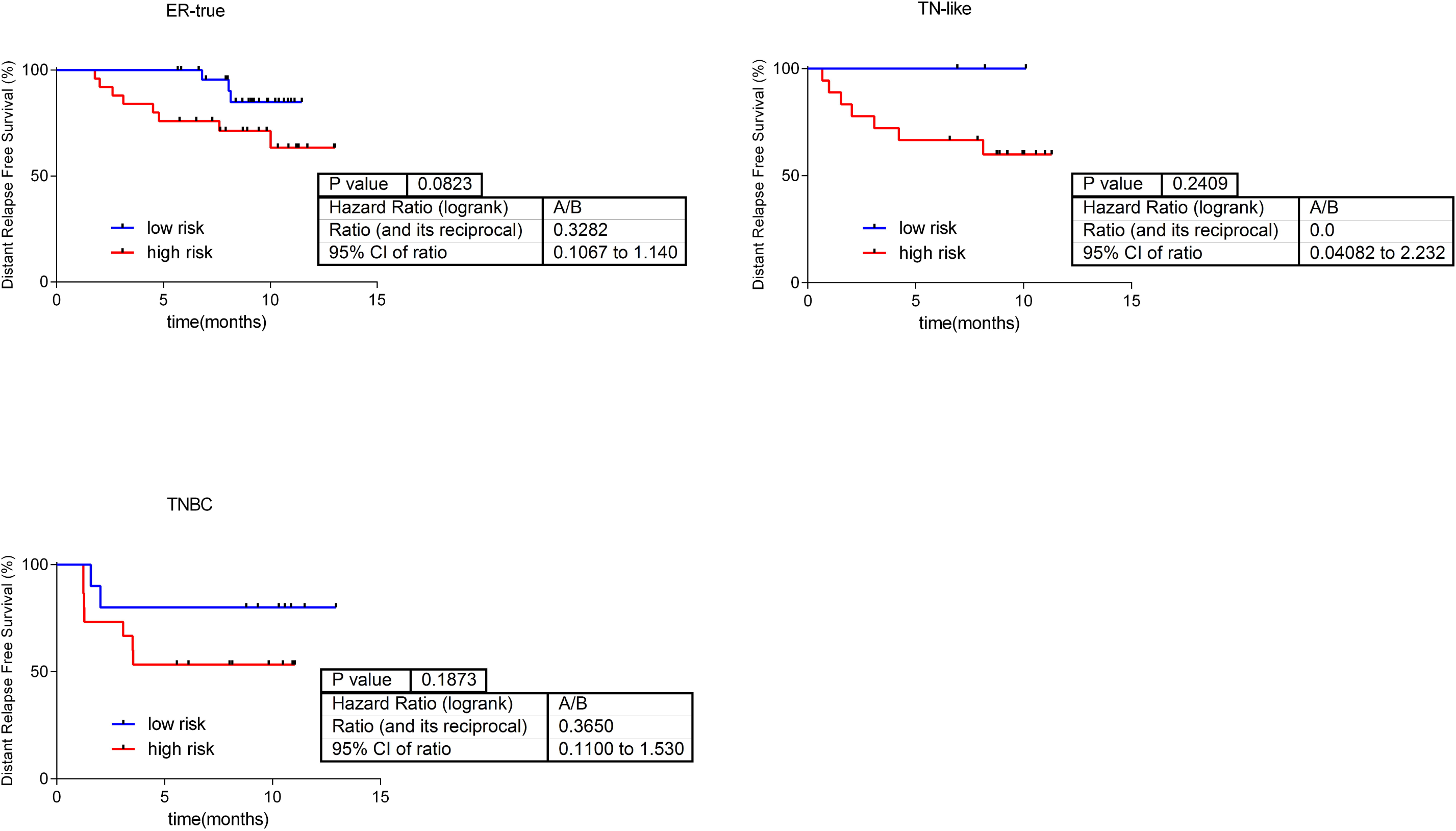
Component 10b activity prognostic value by subtype.

Component 10b contains 10 proteins. Four out of nine edges ( DYNLRB1-HSPH1; HSPH1-CFL1; HBB-UCHL5; and UCHL5-CFL1) have been previously described in G2F database [7]. Then, G2F added two more proteins (HSPH1 and UCHL5) to this component. Most of the proteins included in this component are related with metastasis processes [8–12], whereas PBDC1 doesn't have any associated function. On the other hand, HIST2H2BF is a histone and HIST2H3PS2 is a histone pseudogene showing a directed relationship (Fig 4).

**Fig 4:**
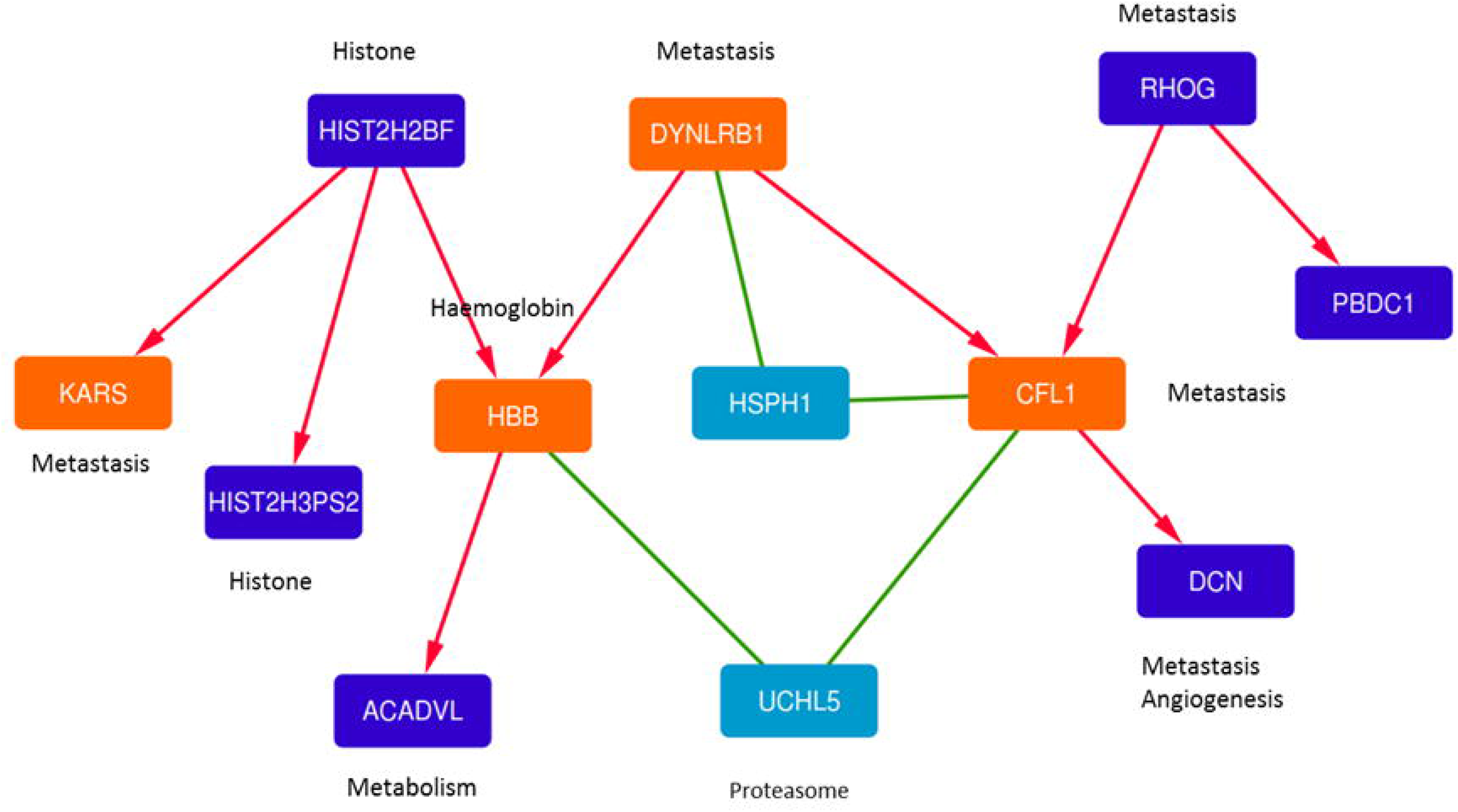
Component 10b merged with PPI information provided by Genes2FANS.

Red arrows indicate relationships from DAG; green lines indicate relationships from Genes2FANS database. Orange nodes are common nodes between these two approaches, purple nodes are nodes which come only from DAG and blue nodes are only from Genes2FANS.

## Discussion

In this study, we used proteomics and DAG to characterize relationships between proteins in breast cancer tumor samples. Unlike other approaches, such as G2F [7], our DAG method supplies directed relationships between proteins and a hierarchical structure. Traditionally, PPI networks are based in relationships described in the literature. However, we built a directed network, i.e. a graph formed by edges with a direction, using protein expression data without other *a priori* information, so it was possible to propose new hypotheses about protein interactions. We used probabilistic graphical models (PGM) because they offer a way to relate many random variables with a complex dependency structure.

Arrows in directed networks indicate causality between two proteins, i.e. if protein A and B are connected and protein A changes its expression value, protein B changes its expression value as well. This approach allows making hypotheses about causal relationships between proteins and proposes a hierarchical structure. G2F relationships supplies additional information to our directed networks, so it served as a validation of the network coherence, i.e., in some cases, an experimental relationship between two proteins connected in the directed network had been previously described. In component 10b, for instance, CFL1 and DYNRBL1 had a described common nexus in HSPH1 [7]. It is remarkable that, in component 27, DAG analysis established a relationship between PTRF and PRDXCDBP which is widely described [13].

We demonstrated in previous works that non-directed graphs provided functional information [2–5]. Interestingly, a functional structure also appeared in this type of networks. Component activities suggested differences in functions such as metastasis, proliferation, proteasome or glycolysis. We have previously described differences in proteasome and glycolysis between ER-true and TN-like subtypes using non-directed networks [2].

On the other hand, it is widely known the role that actin-cytoskeleton plays and its regulation in directional migration and metastasis, Therefore, it is not surprising that the proteins that make up component 10b had some relation to this biological function and had a prognostic value. Using component 10b activity, based on these proteins related with metastasis, it is possible to split our population into a low and a high risk of relapse groups and, interestingly, this prognostic value is also showed in the analysis by subtype. In previous studies we have used functional node activities from non-directed network to develop prognostic predictors [4, 5]. Now, this approach is also validated in directed networks.

Component 10b presents three proteins showing influence over others: HIST2H2BF, DYNLRB1 and RHOG. Dynein light chain roadblock-type 1 (DYNRBL1), also known as km23-1, is a component of the cytoplasmic dynein 1 complex. In colon cells, its depletion can block events known to be involved in cell invasion and tumor metastasis, and it is also an actin cytoskeletal linker critical for the dynamic regulation of cell motility and invasion since it induces a highly organized actin stress fiber network [14]. DYNRBL1 regulates RhoA and motility-associated actin modulating proteins, such as cofilin1 (CFL1) and coronin [15], suggesting that DYNRBL1 may represent a novel target for anti-metastatic therapy [9].

On the other hand, Ras-homolog family member G (RhoG) encodes a member of the Rho family of small GTPases. Constitutively active RhoG induces morphological and cytoskeletal changes similar to those induced by the combination of active Rac and Cdc42 working upstream of them. Rac is activated at the leading edge of motile cells and induces the formation of actin-rich lamellipodia protrusions, which serves as a major driving force of cancer cell. The major downstream proteins for Rac are the WAVE family proteins, the activators of the Arp2/3 complex. In addition, RhoG induces translocation of Dock4-ELMO complex to the plasma membrane and enhances Rac1 activation which promotes migration [16]. Moreover, RNA interference-mediated knockdown of RhoG in HeLa cells reduced cell migration in Transwell and scratch-wound migration assays [11].

HIST2H2BF has been proposed as a pancreatic ductal adenocarcinoma biomarker[17], but there is no available information about its role in metastasis or breast cancer.

In component 10b, both DYNLRB1 and RHOG expression modulates cofilin1 (CFL1). CFL1 is a widely distributed intracellular actin-modulating protein that binds and depolymerizes filamentous F-actin and inhibits the polymerization of monomeric G-actin in a pH-dependent manner. Increased phosphorylation of this protein by LIM kinase aids in Rho-induced reorganization of the actin cytoskeleton [18]. CFL1 is an actin-severing protein that creates free barbed ends in the actin-severing process. Arp2/3 binding to these barbed ends allows the elongation of new actin filaments. This synergy between CFL1 severing activity and Arp2/3-generated dendritic nucleation resulting in the formation of stable invadopods and a directional migration, linking CFL1and ARP2/3, through RhoG, to tumor cell invasion [15, 19]. CFL1 pathway was previously related with metastases processes in breast cancer [18]. Furthermore, DYNRBL1 is required for TGFb1 secretion. It seems to be TGFb pathway plays an important role in cell migration in this sense, because TGFbRII signaling regulates Rho GTPase degradation and actin dynamics. Moreover, TGFb pathway induces expression of the GEF NET1 which accumulates and promotes actin polymerization and on the other hand, it induces expression of tropomyosins that regulate assembly of a contractile apparatus that serves the motility of the responding cell during the EMT process [15] Additionally, TGFß production by macrophages or dendritic cells (DCs) that have engulfed apoptotic cells can promote the generation of inducible regulatory T (Tregs) that play a known protumoral role [16].

Therefore, the relationships between CFL1, RHOG and DYNLRB1 are well-established. Interestingly, DAG graph adds the decorin (DCN) to these edges. DCN is a small leucine-rich proteoglycan that promotes matrix organization by decreasing collagen uptake/degradation, which constitutes a physical barrier against migration/motility of cancerous cells. The alteration of the matrix stiffness leads to differential integrin activation and changes in cytoskeletal organization by Rac, affecting cell motility and invasiveness [20]. Moreover, DCN acts as a matrikine whose interaction with CXC chemokine receptor 4 (CXCR4) impairs its bind with stromal cell-derived factor-1a (SDF-1a), preventing directional migration [20]. In breast cancer tumors, high DCN expression in stroma correlated with lower tumor grade, low Ki67 levels and ER positivity. On the other hand, high expression of DCN in the malignant epithelium was correlated with LN positivity, higher number of positive lymph nodes and HER2-positive status compared to patients with low DCN expression [21].

By last, lysyl-tRNA synthetase (KARS), also known by KRS, was recently shown to induce cancer cell migration through its interaction with the 67-kDa laminin receptor (67LR) who binds laminin on ECM. The interaction of KRS with 67LR enhances the membrane stability of 67LR, which in turn results in an increase laminin-dependent cell migration in metastasis [18]. KARS, causes incomplete epithelial-mesenchymal transition and ineffective cell-extracellular matrix adhesion for migration [22].

We used a mathematical method as DAG analysis and applied them to proteomics data of breast cancer tumors in order to infer causal relationships between these proteins. This method supplied some known relationships but also proposed new ones. Additionally, it associated proteins with a similar function. Therefore, it seems that it is a good approach to propose new hypotheses about mechanisms of action. Moreover, it was possible to associate the results obtained by DAG analysis with prognosis and built a prognostic signature. As far we know, this is the first time that this type of analysis is applied to clinical data and is associated with clinical outcome.

Our study has some limitations. Proteomics provides complementary information to other techniques such as genomics. However, an improvement in the number of detected proteins is still necessary. Validation at the cellular level is also needed.

To sum up, in this study, we used proteomics and directed networks to characterize relationships between proteins in breast cancer tumors. This approach reflected some previously described interactions and it could be used to propose new hypotheses and mechanisms of action.

## Material and methods

### Ethics statement

Informed consent had been obtained for the participants on the study. The approval of the study was obtained by Hospital Doce de Octubre and Hospital Universitario La Paz Ethical Committees.

### Samples

One hundred and six FFPE samples from patients with breast cancer were recovered from I+12 Biobank and from IdiPAZ Biobank, both integrated in the Spanish Hospital Biobank Network. The histopathological characteristics were reviewed by a pathologist to confirm tumor content. Samples had to include no less than half of tumor cells. The endorsement of the study was obtained by Hospital Doce de Octubre and Hospital Universitario La Paz Ethical Committees. These samples were utilized in previous studies [2, 3, 6].

### Protein preparation

Proteins were extracted from FFPE samples as previously described [23]. Briefly, FFPE sections were deparaffinized in xylene and washed twice with absolute ethanol. Protein extracts from FFPE samples were set up in 2% SDS buffer using a protocol based on heat-induced antigen retrieval. Protein concentration was quantified using the MicroBCA Protein Assay Kit (Pierce-Thermo Scientific). Protein extracts (10 μg) were processed with trypsin (1:50) and SDS was removed from digested lysates using Detergent Removal Spin Columns (Pierce). Peptide samples were additionally desalted using ZipTips (Millipore), dried, and resolubilized in 15 μL of a 0.1% formic acid and 3% acetonitrile solution before MS experiments.

### Label-free proteomics

Samples were analyzed on a LTQ-Orbitrap Velos hybrid mass spectrometer (Thermo Fischer Scientific, Bremen, Germany) coupled to NanoLC-Ultra system (Eksigent Technologies, Dublin, CA, USA) as described previously [2, 3]. Briefly, after separation, peptides were eluted with a gradient of 5 to 30% acetonitrile in 95 minutes. The mass spectrometer was operated in data-dependent mode (DDA), followed by CID (collision-induced dissociation) fragmentation on the twenty most intense signals per cycle. The acquired raw MS data were processed by MaxQuant (version 1.2.7.4) [24], followed by protein identification using the integrated Andromeda search engine [25]. Briefly, spectra were searched against a forward UniProtKB/Swiss-Prot database for human, concatenated to a reversed decoyed fasta database (NCBI taxonomy ID 9606, release date 2011-12-13). The maximum false discovery rate (FDR) was set to 0.01 for peptides and 0.05 for proteins. Label free quantification was calculated on the basis of the normalized intensities (LFQ intensity). Quantifiable proteins were defined as those detected in at least 75% of samples in at least one type of sample (either ER+ or TNBC samples) showing two or more unique peptides. Only quantifiable proteins were considered for subsequent analyses. Protein expression data were log2 transformed and missing values were replaced using data imputation for label-free data, as explained in [26], using default values. Finally, protein expression values were z-score transformed. Batch effects were estimated and corrected using ComBat [27]. All the mass spectrometry raw data files acquired in this study may be downloaded from Chorus (http://chorusproject.org) under the project name Breast Cancer Proteomics.

### Network construction

PGM are graph-based representations of joint probability distributions where nodes represent random variables and edges (directed or undirected) represent stochastic dependencies among the variables. In particular, we have used a type of PGM called Bayesian networks (BN) [28]. With these models, the dependences between the variables in our data are specified by a directed acyclic graph (DAG). Firstly, we find the BN that best explains our data [29]. There are different algorithms to learn a DAG from data but we have selected the well-known PC algorithm, a constraint-based structure learning algorithm [30] based on conditional independence tests. The PC algorithm was shown to be consistent in some high-dimensional settings [31] and have some other statistical properties which make it very useful. In this way, our data are represented by a large graph that can be partitioned into several connected components. Then, we focused on finding suitable subgraphs that give us a much more clear understanding of the interrelations therein. All these procedures are implemented in R [32] within packages *pcalg* [31] and *graph* [33]. We used protein expression data without other *a priori* information.

With the aim of compare and complete the information provided by BN, the tool Genes2FANS (G2F), developed by Ma'ayan's group was used [7]. This software provides information about protein-protein interactions (PPI) based on experimental evidence. A PPI network using G2F was built for each component and finally, both networks, the DAG and the PPI network, were merged.

### Gene Ontology Analyses

Protein to Gene Symbol conversion was performed using Uniprot (www.uniprot.org) and DAVID (www.david.ncifcrf.gov) [34]. Gene Ontology Analysis was also done in DAVID selecting only *"Homo sapiens”* background and GOTERM-FAT, Biocarta and KEGG databases. As concerns small connected components, a search in the literature was done so as to establish component main functions.

### Component activity measurements

Component activities were calculated as previously described [3, 4]. Briefly, activity measurement was calculated by the mean expression of all the proteins of each component related with the established major component function.

### Statistical analyses

Network analyses were performed using Cytoscape software [35]. Statistical analyses were done in GraphPad Prism v6. Prognostic signatures were developed using R v3.2.4 and BRB Array Tools, developed by Richard Simon and BRB Array Tools Development Team [36]. P-values under 0.05 were considered statistically significant.

## Funding statement

This study was supported by Instituto de Salud Carlos III, Spanish Economy and Competitiveness Ministry, Spain and co-funded by the FEDER program, “Una forma de hacer Europa” (PI15/01310). LT-F is supported by the Spanish Economy and Competitiveness Ministry (DI-15-07614). GP-V is supported by Conserjería de Educación, Juventud y Deporte of Comunidad de Madrid (IND2017/BMD7783). The funders had no role in the study design, data collection and analysis, decision to publish or preparation of the manuscript.

## Author contributions

All the authors have directly participated in the preparation of this manuscript and have approved the final version submitted. RL-V prepared the proteomics samples. JMA, MD-A, HN and PM contributed the directed graphical models. LT-F, AZ-M, AG-P, G-PV and MF-G performed the statistical analyses, the directed graphical model interpretation and the review of the literature. LT-F, AG-P, JAFV, and EE conceived of the study and participated in its design and interpretation. LT-F and AZ-M drafted the manuscript. AG-P, JAFV, and EE supported the manuscript drafting. AG-P and JAFV coordinated the study.

## Competing interests

JAFV, EE and AG-P are shareholders in Biomedica Molecular Medicine SL. LT-F and GP-V are employees of Biomedica Molecular Medicine SL. The other authors declare no competing interests.

## Data Availability Statement

All the mass spectrometry raw data files acquired in this study may be downloaded from Chorus (http://chorusproject.org) under the project name Breast Cancer Proteomics.

## Supporting information captions

**Sup Table 1: Clinical characteristics of patient cohort.**

**Sup File 1: Components defined by DAG analysis.**

